# An Atypical RNA Quadruplex Marks RNAs as Vectors for Gene Silencing

**DOI:** 10.1101/2021.12.21.472683

**Authors:** Saeed Roschdi, Jenny Yan, Yuichiro Nomura, Cristian A. Escobar, Riley J. Petersen, Craig A. Bingman, Marco Tonelli, Rahul Vivek, Eric J. Montemayor, Marv Wickens, Scott G. Kennedy, Samuel E. Butcher

**Affiliations:** Department of Biochemistry, University of Wisconsin-Madison, Madison, WI, USA; Department of Genetics, Blavatnik Institute, Harvard Medical School, Boston, MA, USA

## Abstract

The addition of poly(UG) (“pUG”) repeats to 3′ termini of mRNAs drives gene silencing and trans-generational epigenetic inheritance in the metazoan *C. elegans*.^1^ pUG tails promote silencing by recruiting an RNA-dependent RNA Polymerase (RdRP) that synthesizes small interfering (si)RNAs.^1^ Here we show that active pUG tails require a minimum of 11.5 repeats and adopt a quadruplex (G4)^2^ structure we term the pUG fold. The pUG fold differs from known G4s in that it has a left-handed backbone similar to Z-RNA^3,4^, no consecutive guanosines in its sequence, and three G quartets and one U quartet stacked non-sequentially. Its biological importance is emphasized by our observations that porphyrin molecules bind to the pUG fold and inhibit both gene silencing and binding of RdRP. Moreover, specific N7-deaza RNA substitutions that do not adopt the pUG fold neither bind RdRP nor induce RNA silencing. These data define the pUG fold as a previously unrecognized RNA secondary structure motif that drives gene silencing. The pUG fold can also form internally within larger RNA molecules. Approximately 20,000 pUG-fold sequences are found in non-coding regions of human RNAs, suggesting the fold likely has biological roles beyond gene silencing.

## Results

The enzyme RDE-3 is required for RNA interference^5^ and catalyzes the non-templated addition of pUG repeats to RNA 3′ ends (hereafter, pUG tails).^1,6^ RDE-3 preferentially initiates pUG tail synthesis with a G, followed by the addition of perfect UG dinucleotide repeats.^6^ pUG tails containing 14 or more dinucleotide repeats recruit the RdRP RRF-1 to RNAs in the *C. elegans* germline to drive gene silencing, while pUG tails harboring only 8 repeats are inactive.^1^ To understand why pUG tails must contain more than 8 dinucleotide repeats, we sought to precisely define the length requirements of a functional pUG tail *in vivo*, in the nematode *C. elegans*. We prepared pUG RNAs composed of the first 541 nucleotides of the *oma-1* coding sequence followed by different lengths of 3′ pUG tails, initiating with a G. The RNAs were injected into the *C. elegans* germline and scored for their ability to silence an embryonic lethal, gain-of-function (*gf*) allele of the *oma-1* gene.^1^ In this inter-generational assay, animals that fail to silence *oma-1(gf)* exhibit embryonic lethality and animals that silence *oma-1(gf)* develop into adult animals at the restrictive temperature for *oma-1(gf)*. Our analysis showed a transition from inactive to active beginning at 11.5 repeats (hereafter denoted (GU)_11.5_): *oma-1* pUG RNAs with (GU)_10.5_ tails were unable to silence *oma-1(gf)* and (GU)_11.5_ tails were able to silence *oma-1(gf)* (Fig. 1A).

**Figure 1.**
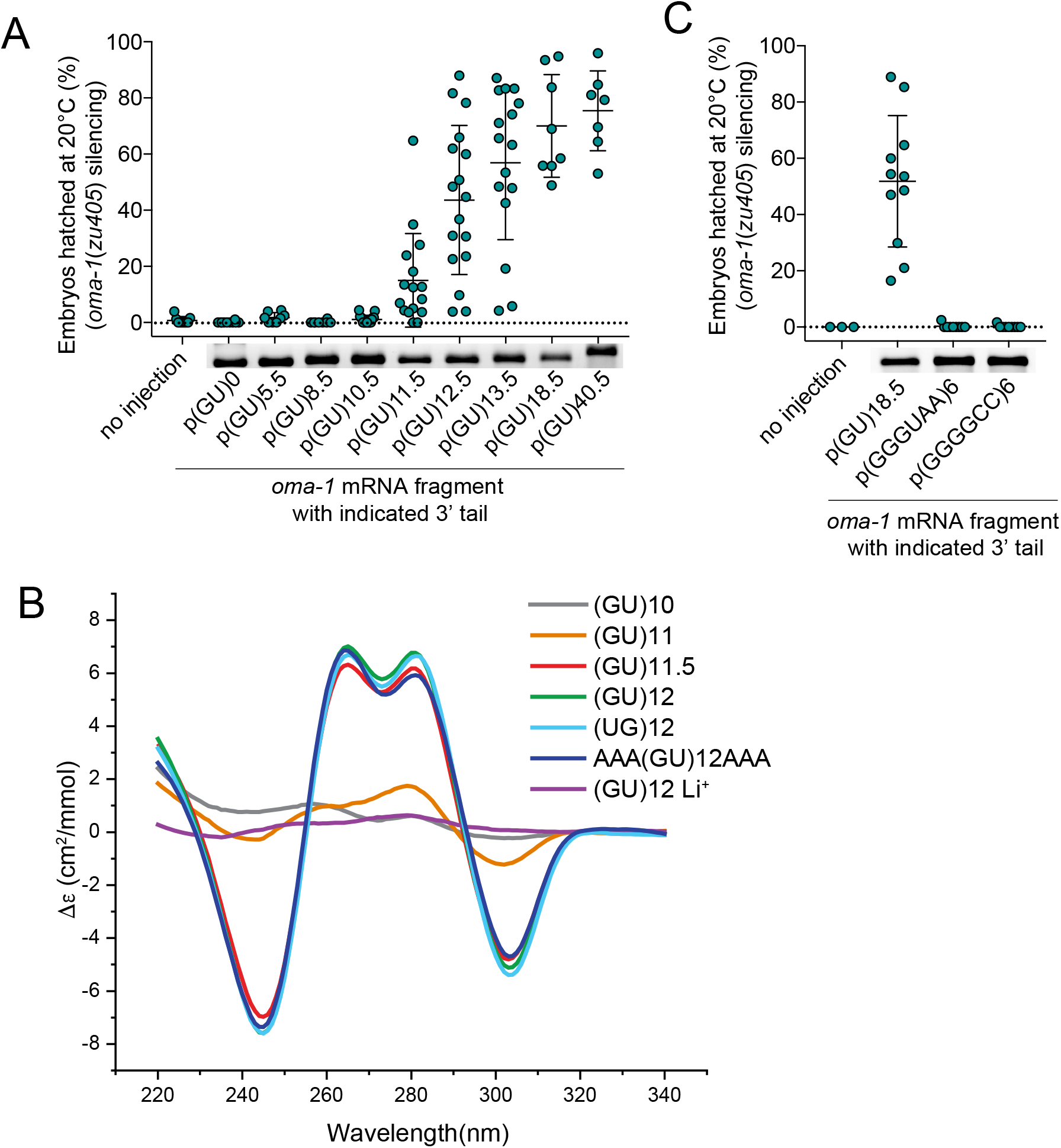
pUG RNA function and folding is length dependent. **a**, RNA silencing in *C. elegans*. oma-1(zu405ts) animals lay arrested embryos at 20°C unless oma-1(zu405ts) is silenced. In vitro-transcribed oma-1 pUG RNAs consisting of the first 541nt of oma-1 mRNA and UG repeat tails of indicated length were injected into the germline of adult rde-1(ne219); oma-1(zu405ts) animal. Each data point represents the percentage of hatched embryos laid by five progeny derived from one injected animal at 20°C (see Methods). Panel below the x-axis shows RNAs run on a 2% agarose gel to assess RNA integrity. Data are mean ± s.d. Number of injected animals, n = 9 (no injection), 10 (p(GU)0), 8 (p(GU)5.5), 8 (p(GU)8.5), 17 (p(GU)10.5), 16 (p(GU)11.5), 17 (p(GU)12.5), 16 (p(GU)13.5), 8 (p(GU)18.5), 7 (p(GU)40.5). To control for possible dsRNA contamination in in vitro transcription reactions, RNAs were injected into rde-1(ne219) mutants, which cannot respond to dsRNA. **b**, RNA secondary structure as a functioned of GU repeat length, measured by circular dichroism spectroscopy. Samples ranged from 10-12 GU repeats. The RNA AAA(GU)12AAA has twelve GU repeats and three adenosines on both the 5′ and 3′ ends. All samples were in 150 mM KCl, except for (GU)12 Li^+^, which was in 150 mM LiCl. **c**, The mammalian telomeric repeat-containing RNA (TERRA) repeat (GGGUAA) and the human C9ORF72 repeat expansion (GGGGCC) consist of hexameric repeats known to adopt G4 structures. *oma-1* mRNA fragments appended with indicated tails were injected into the germlines of adult *rde-1(ne219); oma-1(zu405ts)* animals. Percentage of embryonic arrest of the progeny of injected animals was scored at 20 °C. Data are mean ± s.d. Number of injected animals, n = 6 (no injection), 11 ((GU)18.5), 8 ((GGGUAA)6), and 8 ((GGGGCC)6).

The sharp transition in biological activity at 11.5 GU repeats hinted to us that pUG tails might adopt a secondary structure that is required for their function. To test this hypothesis, we first analyzed the secondary structure of pUG RNAs of various lengths by circular dichroism (CD) spectroscopy. We found that (GU)_11.5_ RNA, but not (GU)_11_ RNA, exhibited a pronounced negative CD absorption peak at 242 nm, a known signature of parallel G quadruplexes (G4s)^7^ (Fig. 1B). G4s are formed when four guanosines engage in Hoogsteen hydrogen bonds (termed a G quartet), which then stack with one or more additional G quartets. G4 structures coordinate potassium ions between quartets, while lithium ions are too small to make such an interaction and destabilize the structures.^8^ The 242 nm CD absorption peak disappeared when (GU)_12_ RNA was prepared in lithium, supporting the idea that the pUG RNA contains a G4 (Fig. 1B). CD analysis also demonstrated that both (GU)_12_ and (UG)_12_ form the same G4 secondary structure (Figure 1B). We find that pUG sequences of 13.5 repeats can tolerate AA substitutions while maintaining silencing function and folding (Supplemental Fig. 1). Additionally, a free 5′ or 3′ terminus is not required as the same fold is observed when adenosine trinucleotides are present on both ends (AAA(GU)_12_AAA, Fig. 1B). Interestingly, the CD spectra also show a negative peak at 304 nm which deviates from the known spectral properties of G4s,^7^ suggesting an unusual structure. In support of this, we found that two known G4 RNAs (UUAGGG)_6_ ^9^ and (GGGGCC)_6_ ^10^, were unable to functionally replace a pUG tail and elicit gene silencing when injected into the *C. elegans* germline (Fig. 1C). Finally, the (GU)_11.5_ structure is stable enough to form under physiological conditions, with a melting temperature of 51.5°C in 150 mM potassium (Supplemental Figure 2). Together, the data suggest that pUG RNA folds into a previously unknown G4 structure (henceforth, “pUG fold”), that requires a minimum of 11.5 GU repeats to form. The striking concordance between the length requirements for *in vivo* function (Fig. 1A), and *in vitro* folding (Fig. 1B), hints that pUG tails may need to adopt the pUG fold to drive gene silencing in the *C. elegans* germline. We therefore sought to define the pUG fold using NMR and crystallography.

**Figure 2.**
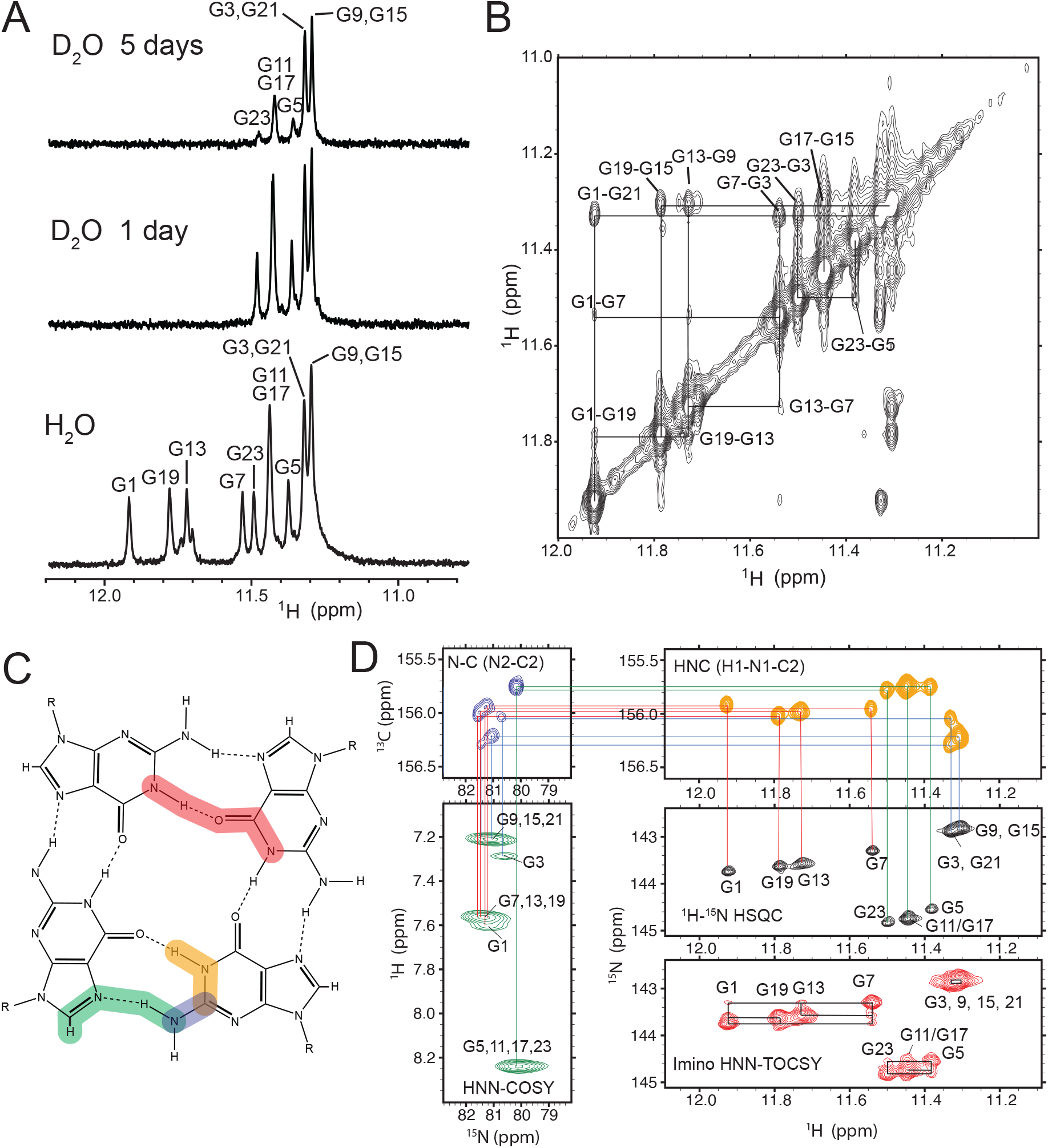
NMR data showing formation of three distinct G quartets in (GU)12. **a**, 1D ^1^H NMR spectra of guanosine imino protons in H_2_O/D_2_O. Bottom spectrum is in H_2_O/D_2_O (90%/10%). Middle and top spectra are 1 day and 5 days after exchange into 99.99% D_2_O, respectively. Resonance assignments are indicated above peaks. **b**, 2D ^1^H-^1^H NOESY spectrum of the imino region showing NOE crosspeaks between guanosines. **c**, Schematic of a G quartet showing through hydrogen bond magnetization transfer pathways. Red shows the HNN-TOCSY pathway, and green shows the HNN-COSY pathway (each pathway occurs four times per quartet but is only diagrammed once for clarity). **d**,Triple resonance ^1^H,^15^N,^13^C experiments were used to connect the through hydrogen bond HNN-COSY correlations to ^1^H-^15^N imino resonances (black) via N2-C2 (blue) and H1-N1-C2 (orange) correlations. HNN-TOCSY through hydrogen bond correlations (red) between the three G quartets are shown with black lines. Spectra are colored as in (c), except for the ^1^H,^15^N HSQC, shown in black.

NMR analysis revealed that most guanosine imino resonances within a pUG fold have dispersed chemical shifts, which is surprising for a simple dinucleotide repeat (Fig. 2A). The guanosine imino resonances display slow rates of hydrogen/deuterium (H/D) exchange, with all but 4 imino resonances observable 5 days after transfer into 99.99% D_2_O, indicating that the pUG fold is kinetically stable at room temperature (Fig. 2A). 2D Nuclear Overhauser Effect spectroscopy (2D NOESY) (Fig. 2B) and through-hydrogen bond correlations (Figure 2C-D) reveal that all twelve guanosines in the pUG fold form three distinct sets of G quartets. For example, all the guanosines form stable amino-N7 and imino-O6 hydrogen bonds (Fig. 2C-D). Substitution of guanosine N7 atoms for carbon (7-deaza G) unfolds the structure (Supplemental Figs. 3A and C). Additionally, the pUG fold is intramolecular as indicated by concentration-independence of electrophoretic mobility on native gels (Supplemental Fig. 3B). Together, the three distinct quartets observed by NMR and the pUG fold’s intramolecular nature explain why 12 Gs are required for structure formation.

**Figure 3.**
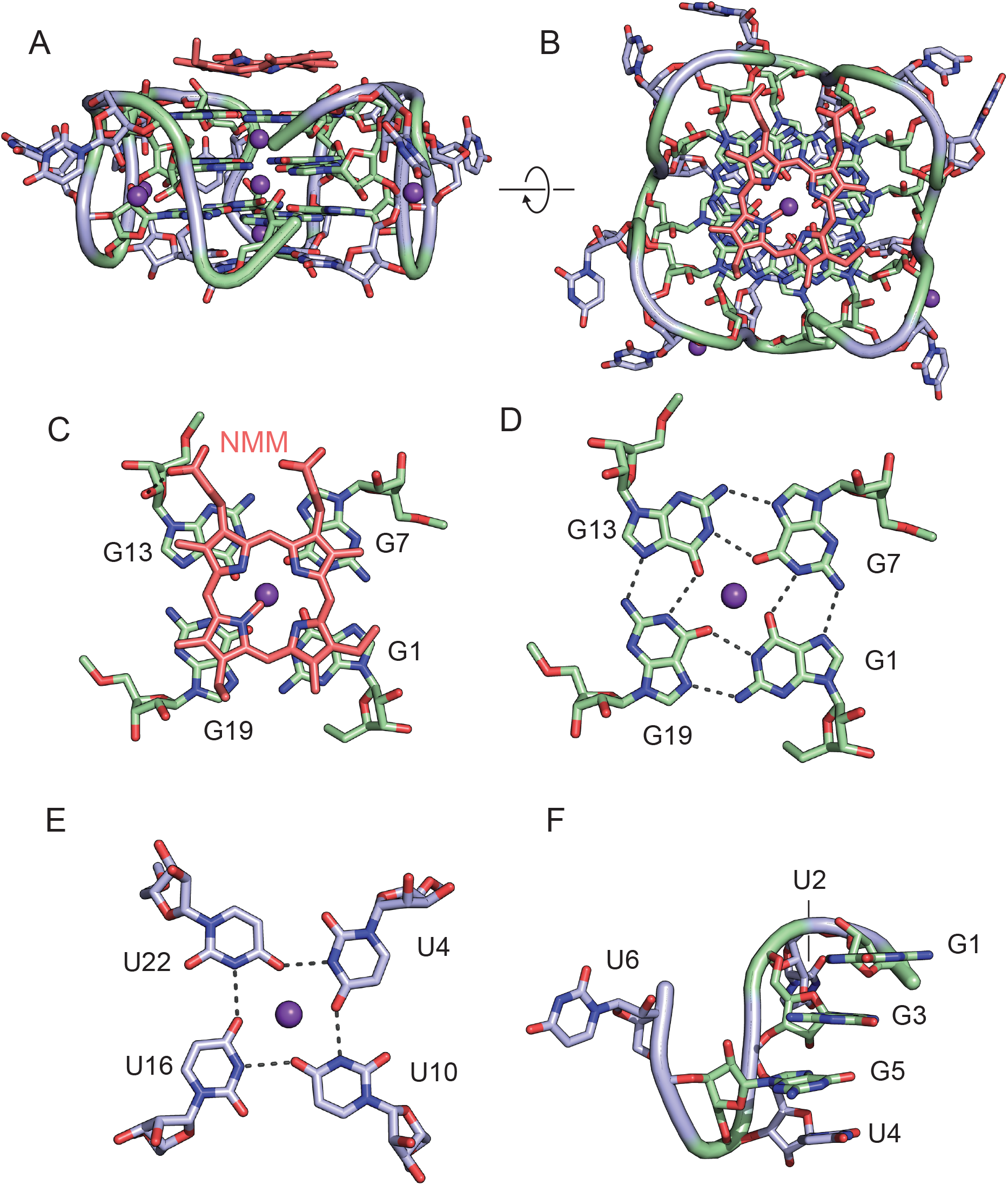
X-ray crystal structure of the (GU)12-NMM complex. **a, b**, Views of (GU)12-NMM, rotated by 90°. Guanosines are green, uridines are blue, potassium ions are purple (shown as one-fourth reduced size for clarity) and NMM is red. The structure of (GU)11.5-NMM is indistinguishable from (GU)12-NMM due to lack of electron density for the 3′ terminal uridine. **c**, Close-up view of NMM interaction with the G1 quartet. A hydrogen bond between a carboxylate oxygen of NMM and the G13 ribose 2′ hydroxyl is shown with a dashed line. **d**, Hydrogen bonds between guanosines in the *syn* G1 quartet. **e**, Hydrogen bonds between the U quartet. **f**, Close-up view of one-fourth of the molecule showing 1-3-5-4 stacking order and backbone turn, a conformation that is repeated four times in (GU)12.

N-methyl mesoporphyrin IX (NMM) is a small molecule that binds parallel DNA and RNA G4s.^11^ NMM binds to the pUG fold with a *K*_d_ of 1 μM (Supplemental Fig. 4A-C). We determined the structures of (GU)_12_ and (GU)_11.5_ in the presence of NMM by X-ray crystallography. Phases were determined via single-wavelength anomalous diffraction (SAD) using a heavy atom derivative obtained by soaking crystals in Rb^+^ (Supplemental Tables 1 and 2). Both RNAs yielded the same crystal form and the resulting electron density maps were sufficient to determine the pUG fold structure to a maximum resolution of 1.97 Å (Figure 3, Supplemental Figure 5 and Supplemental movie). No electron density was observed for the last uridine nucleotide in (GU)_12_, suggesting it is disordered. The structure of the pUG-NMM complex reveals an intramolecular parallel G4 with 3 G quartets and one U quartet (Figure 3 and Supplemental Figure 5). NMR 2D NOESY and through hydrogen-bond measurements (Figure 2) and residual dipolar coupling analysis (R^2^=0.96, Supplemental Fig. 6) indicate the structure of free (GU)12 in solution is very similar to the pUG-NMM crystal structure. The structure adopts an overall left-handed^12^ fold with Z-form RNA^3,13^ *syn-anti* stacking interactions (Supplemental Figure 7). The structure makes a full turn, placing the 5′ and 3′ ends only 3.3 Å apart in (GU)_11.5_ (Supplemental movie). Aside from the chemically distinct 5′ and 3′ ends, the structure displays four-fold symmetry, such that the conformation of the first 6 nucleotides is repeated 4 times.

**Figure 4.**
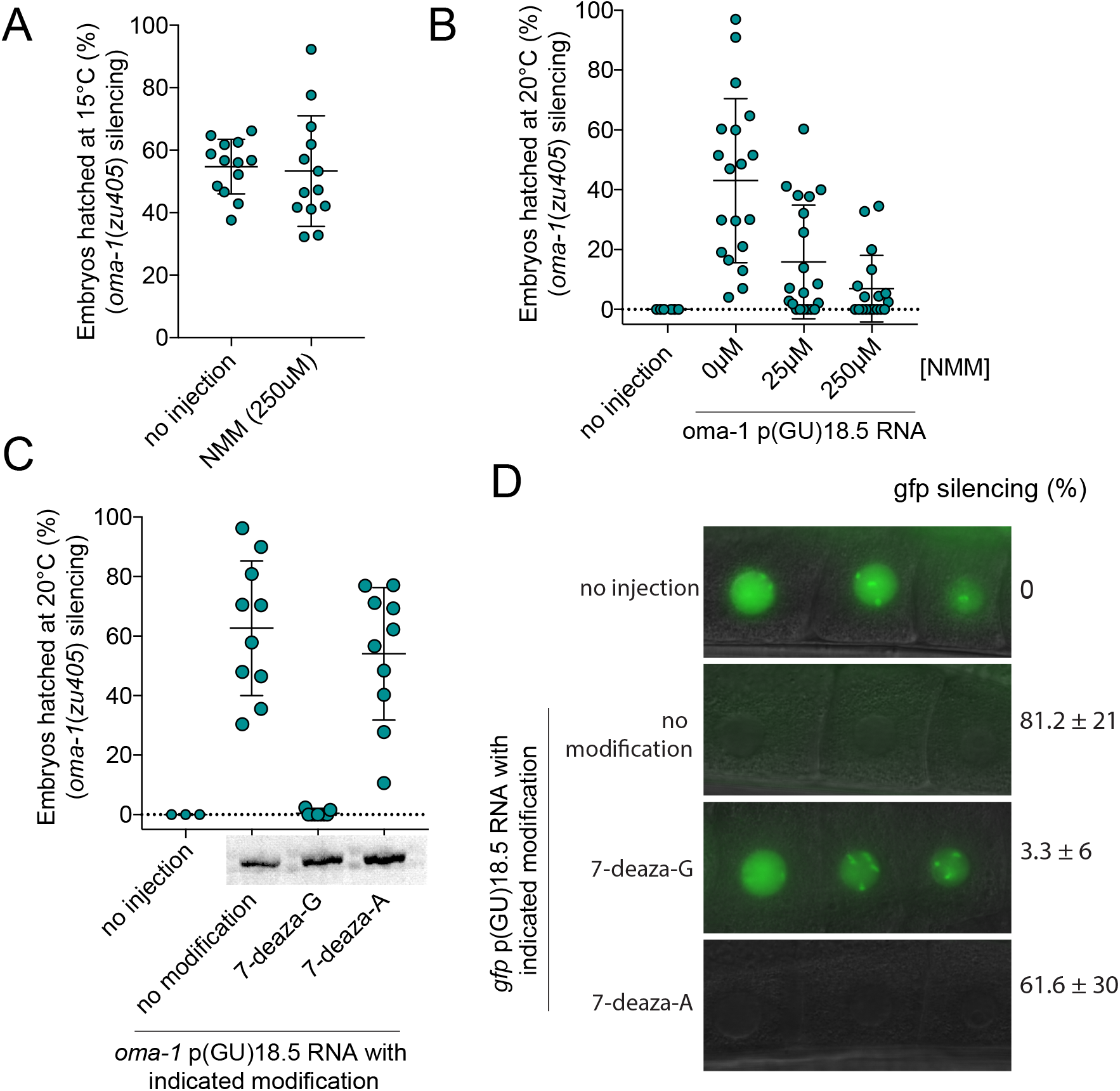
Silencing activity of pUG RNAs is blocked by NMM and substitution with 7-deaza-G. **a**, Injection of 250 μM NMM into the germlines of *rde-1(ne219); oma-1(zu405ts)* animals does not affect embryo hatching at 15°C, showing that NMM does not impact the embryo survivability of *rde-1(ne219); oma-1(zu405ts)* animals. Data are mean ± s.d. Number of injected animals, n = 13 (no injection), and 13 (250μM NMM). **b**, oma-1 pUG RNAs consisting of the first 541nt of oma-1 mRNA and 18.5 GU repeat tails were pretreated with or without indicated concentrations of NMM and injected into the germlines of adult *rde-1(ne219); oma-1(zu405ts)* animals (see Methods). Percentage of embryonic arrest of the progeny of injected animals was scored at 20 °C. Data are mean ± s.d. Number of injected animals, n = 6 (no injection), 19 (no treatment), 20 (25 µM NMM) and 18 (250 μM NMM). **c**, Incorporation of 7-deaza-G blocks the silencing activity of oma-1 pUG RNAs, while control pUG RNAs containing 7-deaza-A substitutions remain functional. Adult rde-1(ne219); oma-1(zu405ts) animals were injected in the germline with oma-1 pUG RNAs consisting of the first 541nt of oma-1 mRNA and 18.5 GU repeat tails. Percentage of embryonic arrest of the progeny of injected animals was scored at 20 °C. Data are mean ± s.d. Panels below the x-axis show RNAs run on 6% polyacrylamide gel to assess RNA integrity. Number of injected animals, n = 3 (no injection), 10 (no modification), 9 (7-deaza-G), and 10 (7-deaza-A). **d**, pUG RNAs synthesized with 7-deaza-G blocked failed to silence gfp, while control pUG RNAs containing 7-deaza-A substitutions remained functional. Percentage of progeny with gfp::h2b silenced (mean ± s.d.). Number of injected animals, n = 5 (no injection), 8 (no modification), 8 (7-deaza-G), and 8 (7-deaza-A). Fluorescence micrographs showing -1, -2, and -3 oocytes of adult progeny of rde-1(ne219); gfp::h2b animals after germline injection of in vitro-transcribed gfp pUG RNAs, consisting of the first 369nt of gfp mRNA and 18.5 GU repeat tails.

**Figure 5.**
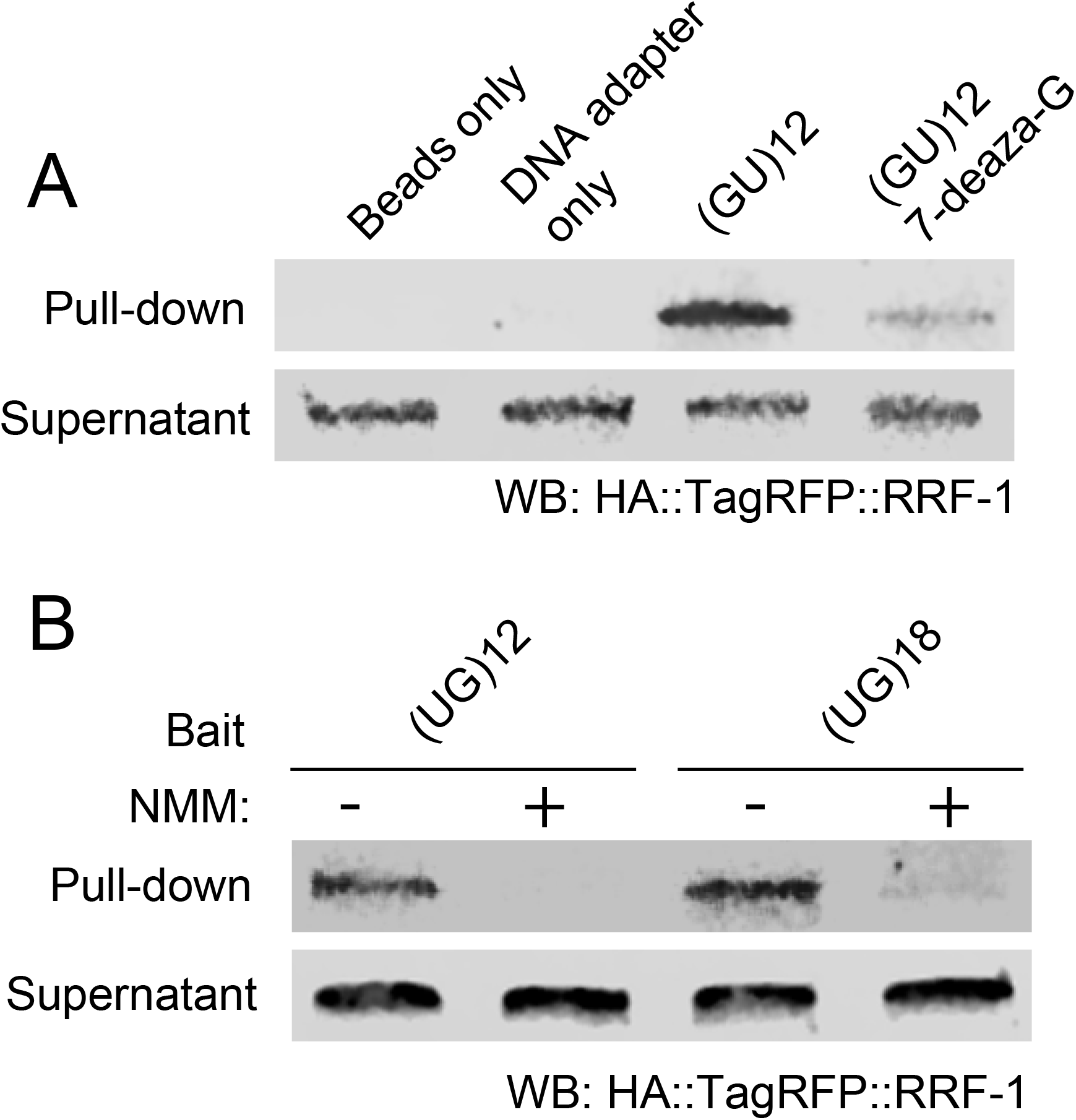
A. 7-deaza-G substitutions and NMM prevent interactions between GU repeats and RRF-1. RNA oligonucleotides consisting of a random 30 nt sequence followed by 12 GU repeats were synthesized with or without 7-deaza-G and conjugated to streptavidin beads using a 3’ biotinylated DNA adapter (see Methods). RNA-conjugated beads were then incubated with extracts from animals expressing HA::TagRFP::RRF-1. Bead-bound material (pull-down) and supernatant were analyzed by Western blot using HA antibodies. Data are representative of two biologically independent experiments. **b**, NMM treatment prevented interactions between GU repeats and RRF-1. 5′ biotinylated RNA oligonucleotides of indicated UG repeat length were conjugated streptavidin beads with or without pretreatment with 250 μM NMM. RNA-conjugated beads were then incubated with extracts from animals expressing HA::TagRFP::RRF-1. Bead-bound material (pull-down) and supernatant were analyzed by immunoblotting with HA antibody. Data are representative of two biologically independent experiments.

Nucleotides G1-G7-G13-G19, (hereafter, the G1 quartet) constitute the first layer of the pUG fold (Fig. 3 A-D) and interact with the porphyrin ring of NMM. The G1 quartet guanosines are in the *syn* conformation and inverted, making this quartet locally right-handed despite the overall left-handed fold (Fig. 3D). After the G1 quartet is a layer of bulged uridines (U2, U8, U14 and U20) (Fig. 3A, B and F). The bulged U2 layer allows the G3 quartet (G3-G9-G15-G21) to stacks below the G1 quartet (Fig. 3F). The G3 quartet is also in the *syn* conformation, but unlike the G1 quartet is in a left-handed orientation. Stacked below the G3-layer quartet is the G5-layer quartet (G5-G11-G17-G23), which is in the *anti* conformation. The left-handed Z-form^3,13^*syn-anti* stacking of the G3 and G5 layer quartets comprise the center of the molecule (Supplemental Fig. 6). Stacked below the G5-layer is a uridine quartet (U4-U10-U16-U22) (Fig. 3E). This U4 layer quartet is in the *anti* conformation. The G5 layer is able to insert between the G3 and U4 layers due to a sharp bend in the backbone following the U4 layer (Fig. 3F). Finally, the U6 layer (U6, U12 and U19) forms three single uridine propeller loops (Fig. 3F). Three potassium ions are coordinated between each of the 4 G/U quartets, and electron density is observed for an additional 3 potassium ions bound to the backbone between the ribose oxygens of the bulged U2 layer and the G3 quartet (Fig. 3A, Supplemental Figure 5A, and Supplemental movie). In summary, the pUG fold is: 1) a G4 with no consecutive guanosines, 2) an overall left-handed G4 with Z-form stacking, 3) comprised of a mixture of G quartet types and one U quartet, and 4) non-sequentially stacked in the order G1-G3-G5-U4.

NMM stacks on top of the G1 quartet in a manner that likely explains the specificity of NMM for parallel RNA G4s^11^ by forming a hydrogen bond between one of its carboxyl groups and the 2’ OH of G13 (Fig. 3A, B and C). Incomplete electron density is also observed for NMM stacking on the U quartet. NMM binding does not alter the pUG fold (Supplemental Fig. 8A) and, in fact, increases the T_m_ from 51.5°C to 59.7°C (Supplemental Fig. 8B). The interaction we observe between NMM and (GU)_12_ RNA suggests that other porphyrins, such as hemin, should also bind to the pUG fold with similar affinity. Indeed, we find that hemin binds (GU)_12_ RNA similarly to NMM (Supplemental Fig. 4 D-F, and see Discussion).

According to current models, pUG tails convert RNAs into agents of gene silencing by recruiting RdRP enzymes that use pUG-tailed RNAs as templates to synthesize gene-silencing small interfering (si)RNAs.^1^ To begin investigating if the pUG fold contributes to pUG RNA function, we asked if RNA sequences capable of adopting a pUG fold (12 or more repeats) might be depleted from *C. elegans* UTRs, as would be expected for an RNA element that silences its host mRNA by recruiting RdRP. The analysis showed that indeed, the *C. elegans* 3’UTRome does contain pUG repeats of less than 12 units but none of 12 or more repeats. In contrast, humans lack an RdRP and have 383 3′ UTRs with 12 or more GU repeats (Supplemental Fig. 9). pUG fold compatible repeats were found in both *C. elegans* and human introns (Supplemental Fig. 9) and were particularly abundant (>19,000) in human introns, where they were non-randomly distributed (see discussion). The genomic distribution of pUG fold elements in the *C. elegans* genome is consistent with a role for this structure in mRNA silencing.

To further probe whether pUG-fold formation is required for gene silencing *in vivo*, we asked whether the pUG ligand NMM affected pUG RNA-based gene silencing. We find that pre-treatment of *oma-1* pUG RNAs with NMM prevented *oma-1* pUG RNA from silencing *oma-1* in a dose-dependent manner (Fig. 4B). Control injections showed that NMM by itself had no observable effect on fertility or development (Fig. 4A). Next, we tested pUG tails containing 7-deaza G modified nucleotides that, based upon native gel analysis and CD spectra (Supplemental Figures 3A and C) do not adopt the pUG fold. We injected 7-deaza-G containing pUG RNAs into the *C. elegans* germline and observed that *oma-1* or *gfp* pUG RNAs with this modification were unable to trigger *oma-1* or *gfp* gene silencing, respectively (Fig. 4C and D). Control *oma-1* and *gfp* pUG RNAs synthesized with unmodified nucleotides or 7-deaza-A modified adenosine retained gene silencing activity (Fig. 4C and D). Thus, targeting the pUG fold, without altering the underlying pUG sequence, disrupts silencing by pUG RNAs. Finally, because pUG tails are thought to recruit RdRP, we asked if 7-deaza-G or NMM might affect the ability of the RdRP RRF-1 to associate with pUG RNAs. Indeed, pUG RNAs containing 7-deaza-G modified guanosine nucleotides failed to associate with RRF-1 (Fig. 5A) and treatment of pUG RNAs with NMM prevented pUG RNAs from interacting with RRF-1 (Fig. 5B). In summary, three lines of evidence show that the pUG fold forms *in vivo* and is required for gene silencing. First, a striking concordance in length requirements for pUG fold formation *in vitro* and gene silencing activity *in vivo* suggests biological function. Second, the genomic profile of pUG fold capable RNA elements is consistent with a role in mRNA silencing. Third, interventions that specifically target the pUG fold structure inhibit RdRP recruitment and gene silencing *in vivo*.

## Discussion

Here we define a previously unrecognized type of RNA secondary structure, the pUG fold, that contributes to gene silencing in *C. elegans* by recruiting RdRP to synthesize siRNAs. RdRP-based gene silencing in *C. elegans* is feed-forward and transgenerational and, therefore, ensuring accuracy within the system is paramount. The exacting sequence requirements and unique secondary structure of the pUG fold is likely one such safe-guard that ensures only pUG-tailed RNAs serve as templates for RdRP. The pUG fold provides a unique platform for reversing the direction of the RNA backbone, consistent with previous models positing pUG tails act as primers for RdRP-based transcription.^1,6^

We find that ≌20,000 pUG fold sequences are transcribed in mammalian introns (Supplemental Figures 9 and 10). Because the pUG fold is tolerant of short insertions (Supplemental Figure 1), the actual number may be even greater. Although robust helicase activities can unfold G4s in spliced mRNAs,^14^ intronic G4s have been less well studied. A growing body of evidence indicates that RNA G4s form in cells and influence gene expression.^15-19^ The pUG fold would not have been identified by existing G4 search algorithms,^20^ implying that G4 RNAs may be more common than currently appreciated. As mammals are not thought to possess an RdRP, the biological function, if any, of mammalian pUG RNAs is unknown. We note, however, that the distribution of pUG fold sequences in the human transcriptome is non-random, with 97% of pUG fold sequences located in introns (Supplemental Fig. 9) where they are enriched near splice sites (Supplemental Fig. 10). Indeed, the splicing regulator TDP-43 (*C. elegans* TDP-1) binds pUG RNA.^1^ Mammalian TDP-43 binds to single stranded pUG RNAs,^21-25^ but has also been observed to bind to G4 RNAs.^26-28^ Condensed TDP-43 aggregates are associated with amyotrophic lateral sclerosis (ALS) and it will therefore be of interest to determine whether pUG RNAs assist in this condensation.^29,30^ In addition to potential roles in splicing, pUG fold sequences are found in lncRNAs^29^ and hundreds of human mRNAs harbor pUG fold sequences within 5′ or 3′ UTR elements, where they may have regulatory function (Supplemental Fig. 9).

pGT/AC is the most common DNA microsatellite repeat in mammalian genomes^31^ and the length of many, if not most, human pGT/AC genomic microsatellite repeats is polymorphic, indicating that the number and identity of RNAs capable of adopting pUG folds is likely to vary substantially in populations and during health and disease. Indeed, expansion/contraction alleles of pGT/AC microsatellite repeats have already been linked to diseases including cystic fibrosis, sterility, asthma, and cancer.^32-42^ In the case of cystic fibrosis, a GU repeat expansion in intron 9 of the CFTR gene from eleven to twelve repeats causes disease,^23,40,41^ and this is precisely the length at which a pUG adopts the pUG fold. These data hint that expansion/contraction of pGT/AC microsatellite repeats can alter the spectrum of RNAs adopting the pUG fold, which could lead to changes in gene expression and, therefore, disease.

We find that hemin binds with high affinity to the pUG fold (Supplemental Fig. 4D-F), which occurs twice in the heme oxygenase-1 (HO-1) pre-mRNA 5′ UTR. HO-1 is rate-limiting for heme catabolism and splicing of HO-1 pre-mRNA is regulated by hemin.^43^ Our data hint at a potential mechanism for this regulation; porphyrin binding to pUG folds can modulate splicing or other RNA-related events. Such a riboswitch-like mechanism would allow cellular levels of HO-1 to be coordinated with porphyrin levels to ensure homeostasis of heme metabolism. Given the abundance of pUG repeat containing RNAs in eukaryotic transcriptomes, and the wide variety of biochemical processes mediated by porphyrins in diverse species, porphyrin-pUG fold interactions might enable small molecule-mediated control of gene expression by design.

## Figure Legends

**Supplemental Figure 1.**
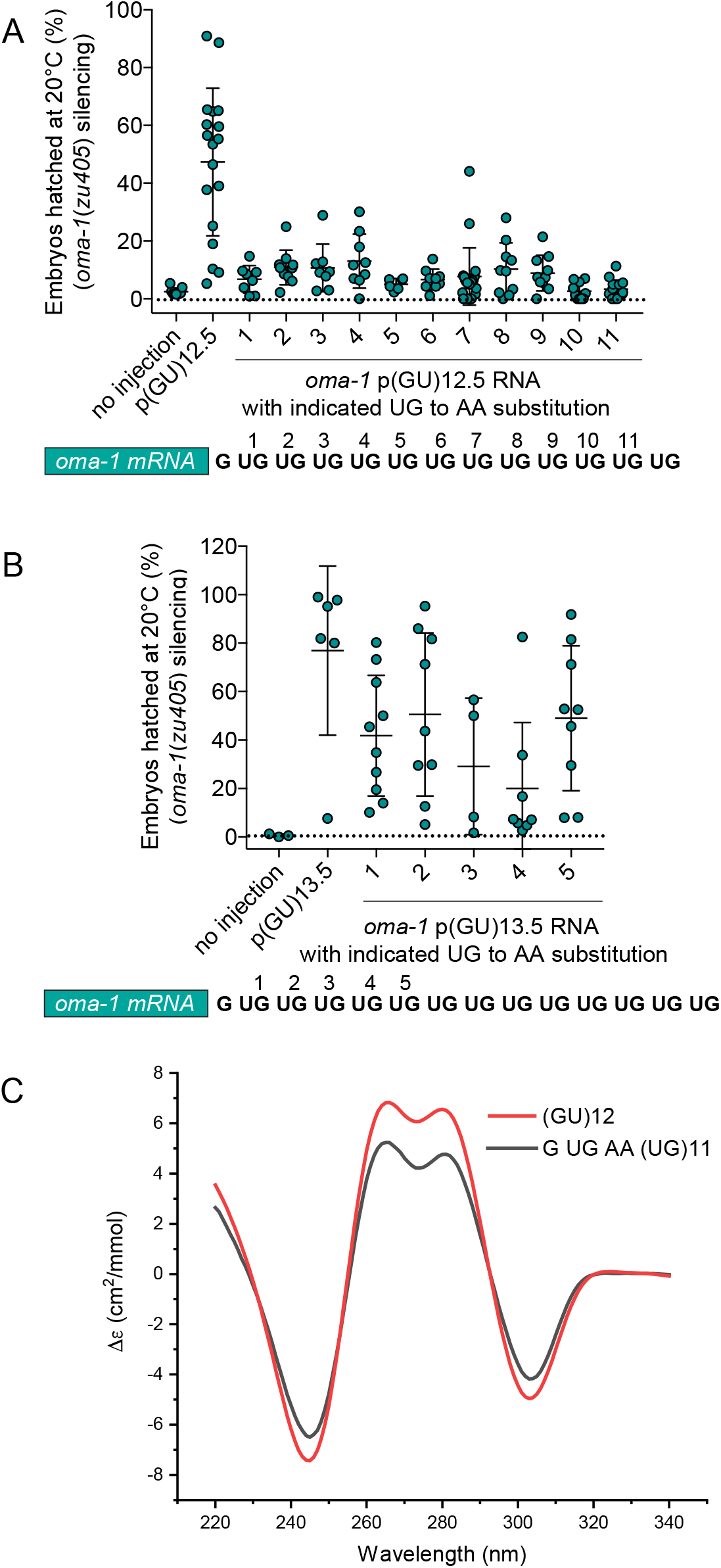
**A**. oma-1(zu405ts) silencing assay with AA substitutions within the pUG tail (GU)12.5. The pUG tail sequence is shown below the plot, with location of AA substitutions indicated at the numbered positions. **B**. oma-1(zu405ts) silencing assay of (GU)13.5 with AA insertions, sequences indicated as in **A. C**. CD secondary structure analysis of (GU)13.5 with AA substitution at position 2, compared to (GU)12.

**Supplemental Figure 2.**
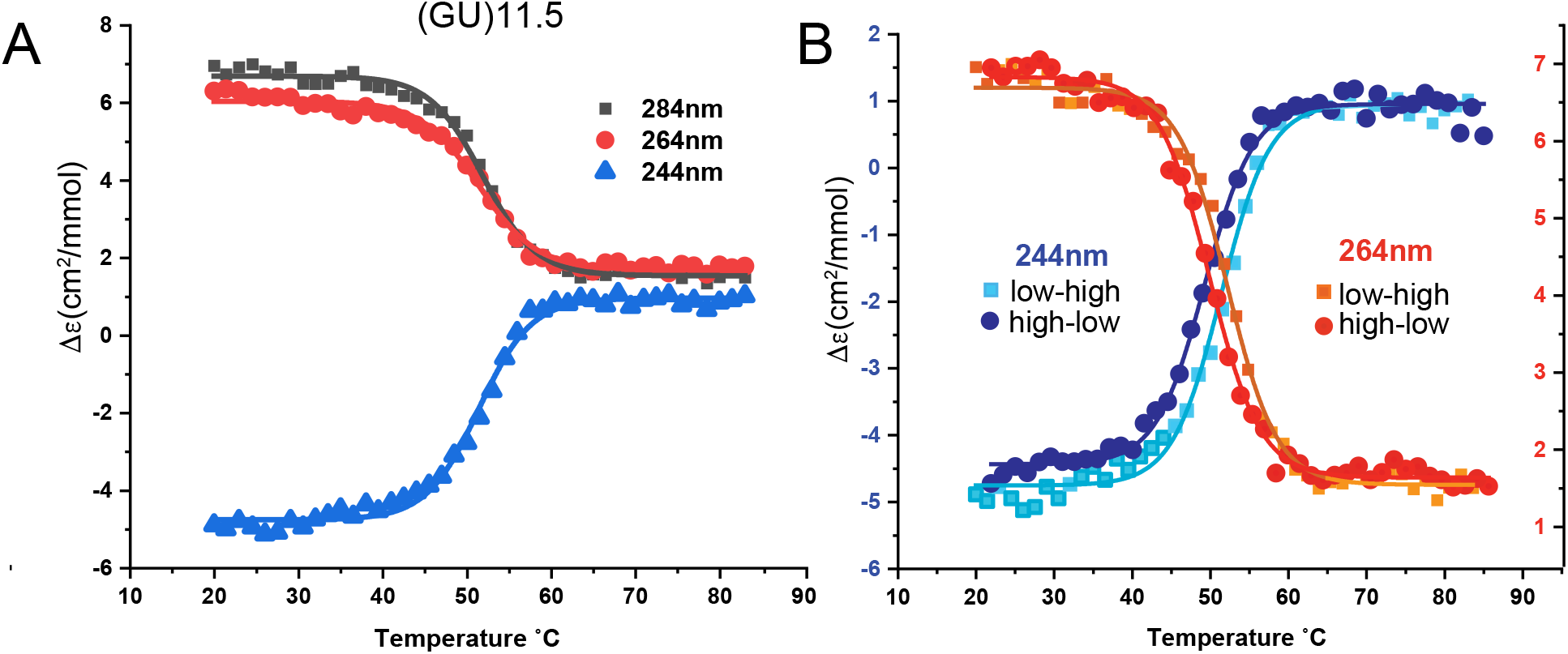
CD monitored thermal denaturation of (GU)11.5 in 150 mM KCl. **A**. Three different wavelengths show a single cooperative melting transition at 51.5 °C. **B**. Thermal melting data measured from low to high temperature and high to low temperature show minimal hysteresis (< 3 °C).

**Supplemental Figure 3.**
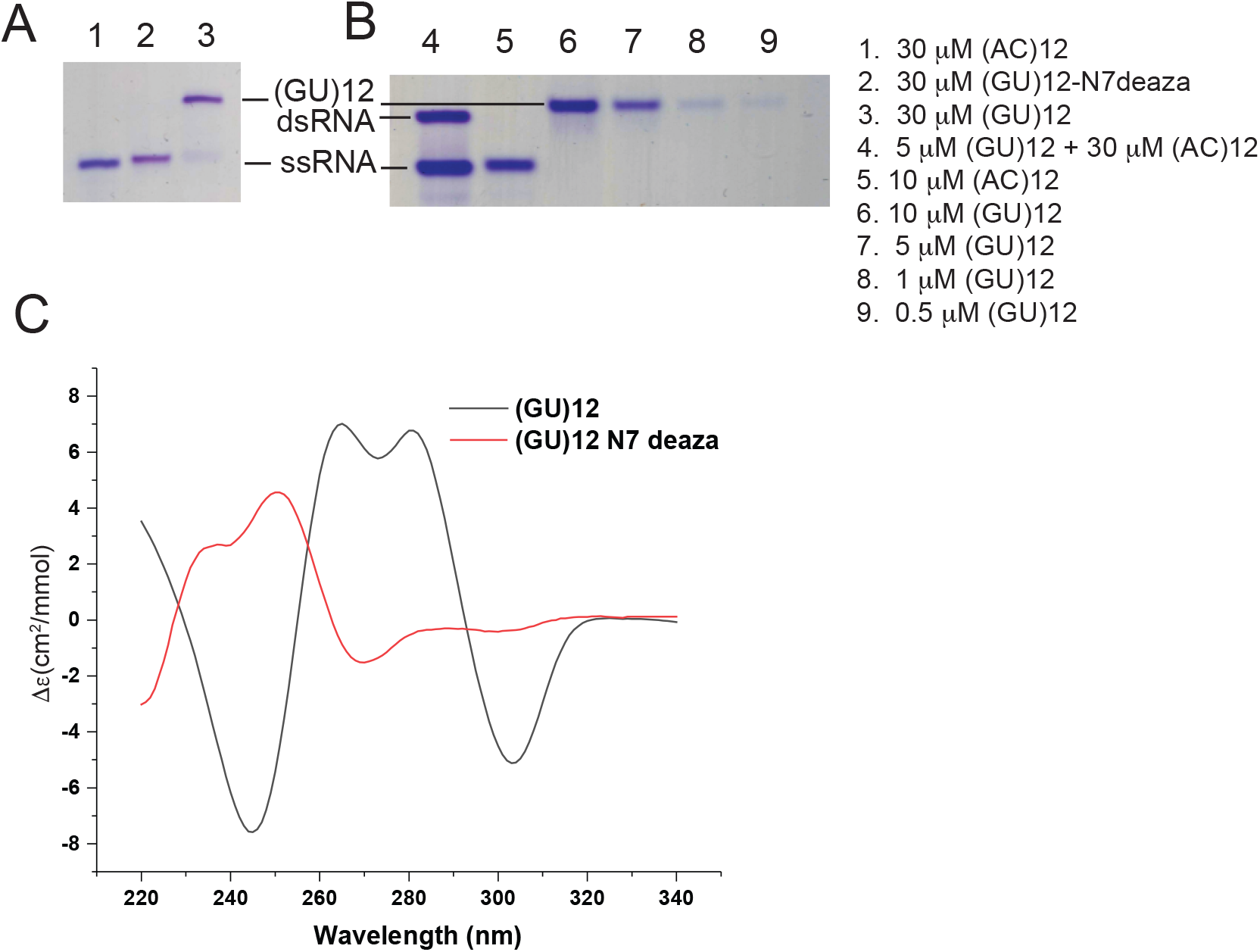
A. pUG RNA is unfolded by N7 deaza G substitution. Native gel analysis of (GU)12 electrophoretic mobility. Lane1: (AC)12 was used as a marker for single stranded RNA (ssRNA). Lane 2: N7 deaza G substitution of (GU)12 produces ssRNA with the same electrophoretic mobility as (AC)12. Lane 3: (GU)12 RNA runs with anomalously slow electrophoretic mobility. B. The pUG fold electrophoretic mobility is concentration independent. Lane 4: double stranded RNA (dsRNA) was enforced by heat annealing (GU)12 to excess (AC)12 complementary ssRNA (lane 2). Lane 5: ssRNA maker (AC)12. Lanes 6-9: (GU)12 at 10, 5, 1, and 0.5 μM, respectively. **C**. CD analysis of unfolded N7 deaza G substituted (GU)12 compared to (GU)12.

**Supplemental Figure 4.**
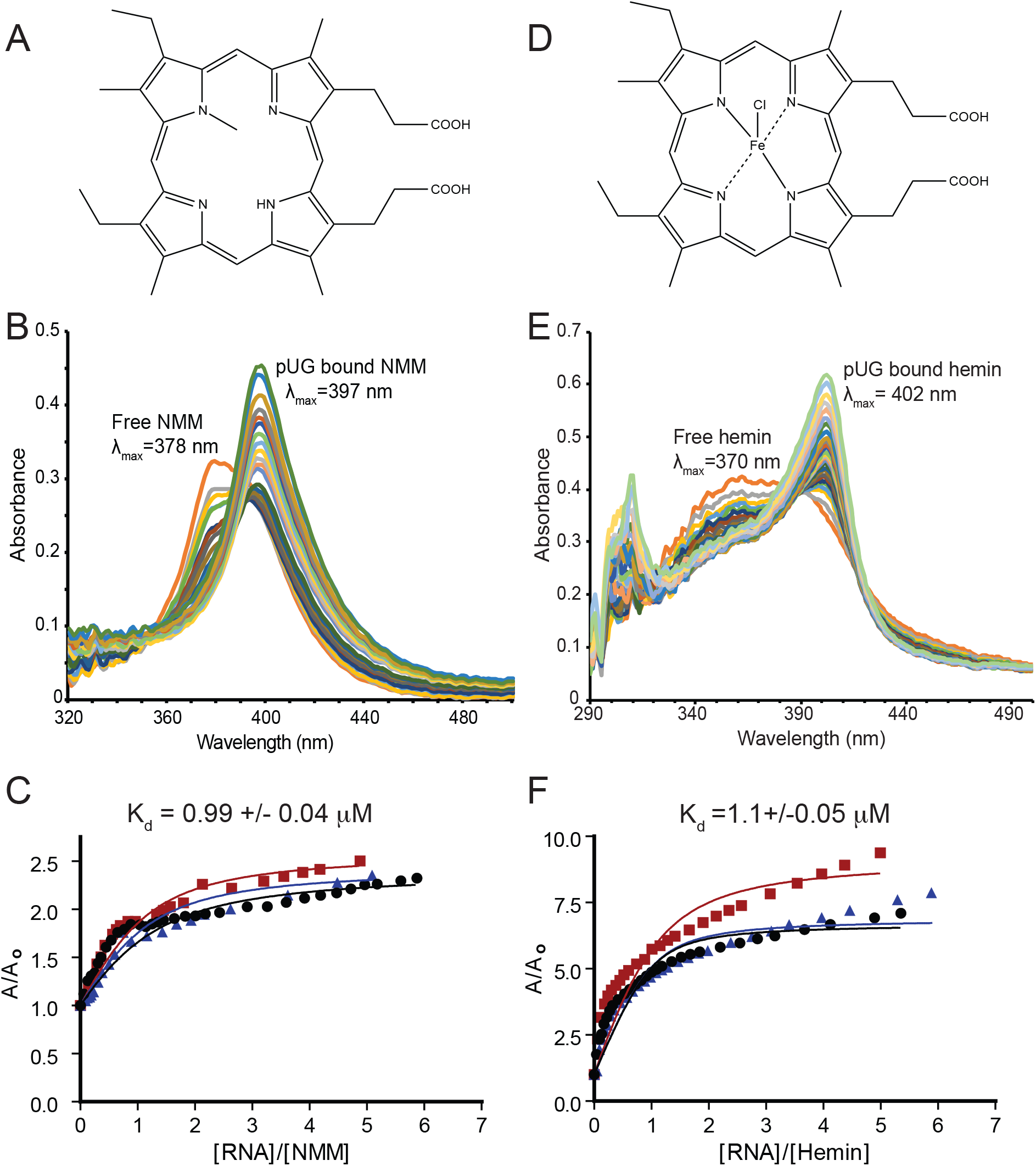
The pUG fold binds the porphyrins NMM and hemin. **A**. Chemical structure of NMM **B**. The NMM absorbance of free NMM (orange, λmax=378 nm) displays a hyperchromic shift to (λmax=397 nM) upon addition of increasing amount of the pUG RNA (GU)11.5. **C**. Fitting of data in A to an equilibrium binding equation. The results of 3 independent experiments are plotted in black, blue and red. **D**. Chemical Structure of hemin **E**. The absorbance of free hemin (orange, λmax=370 nm) displays a hyperchromic shift to (λmax=402 nM) upon addition of increasing amount of the pUG RNA (GU)11.5. **F**. Fitting of data in B to an equilibrium binding equation. The results of 3 independent experiments are plotted in black, blue and red.

**Supplemental Figure 5.**
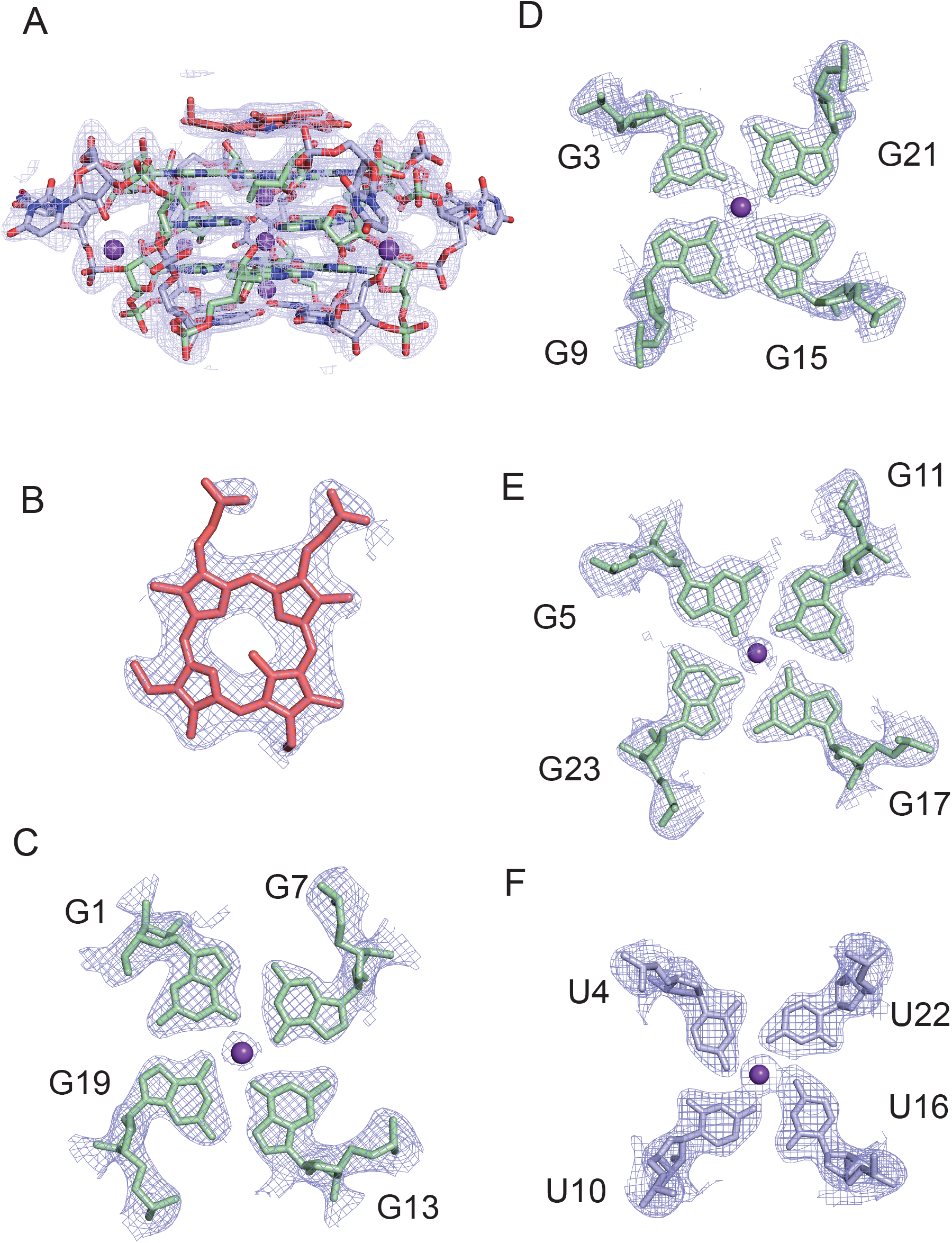
A. Electron density map for (GU)12-NMM contoured at 1 r.m.s.d. B. Electron density for NMM. C. Electron density for the G1 quartet. D. Electron density for the G3 quartet. E. Electron density for the G5 quartet. F. Electron density for the U4 quartet.

**Supplemental Figure 6.**
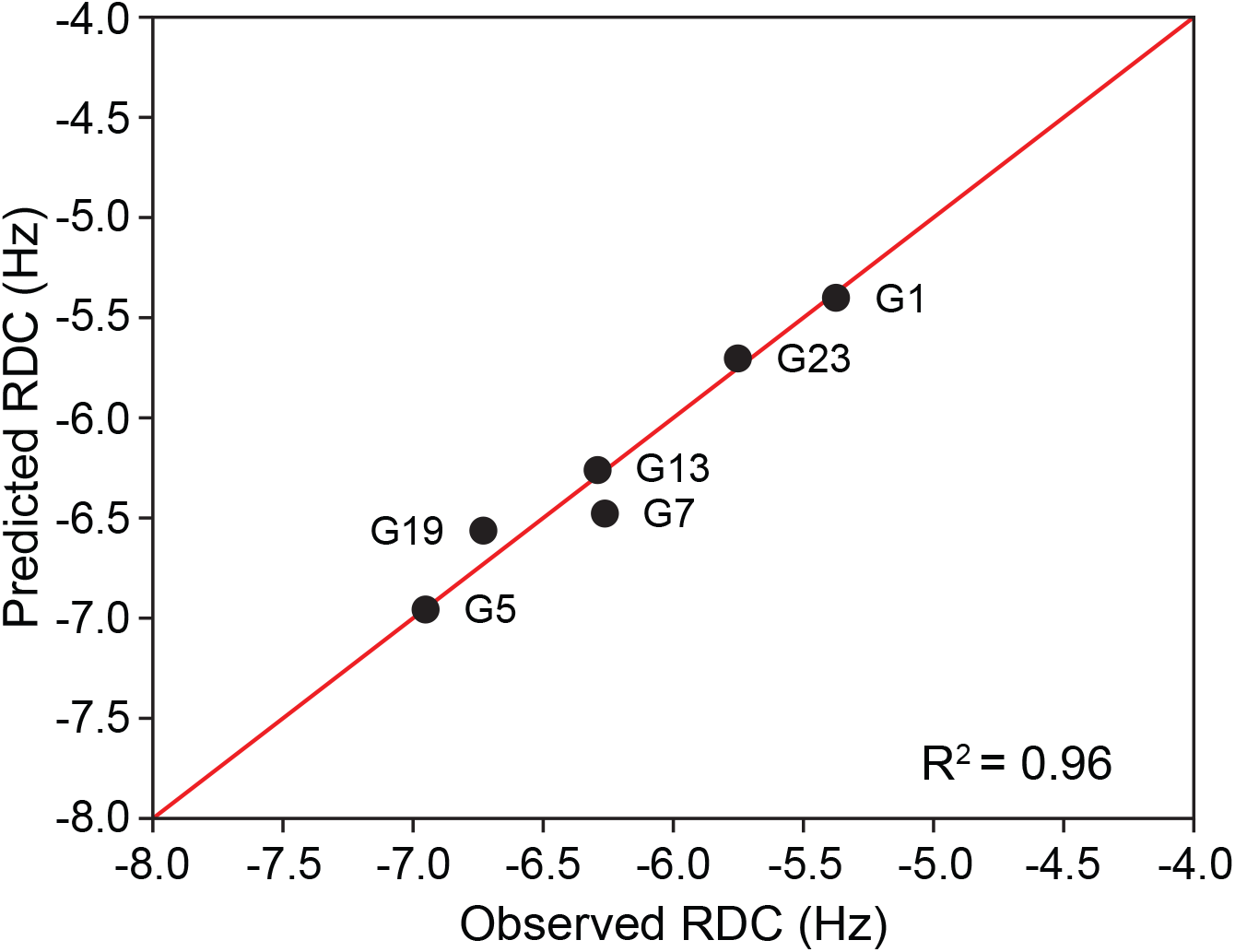
Measured residual dipolar couplings (RDCs) vs. predicted RDCs from the (GU)12-NMM crystal structure. NMR RDCs were measured for ^13^C,^15^N G-labeled (GU)12 RNA (observed) and plotted against the predicted RDC values from the (GU)12-NMM crystal structure, R^2^=0.96.

**Supplemental Figure 7.**
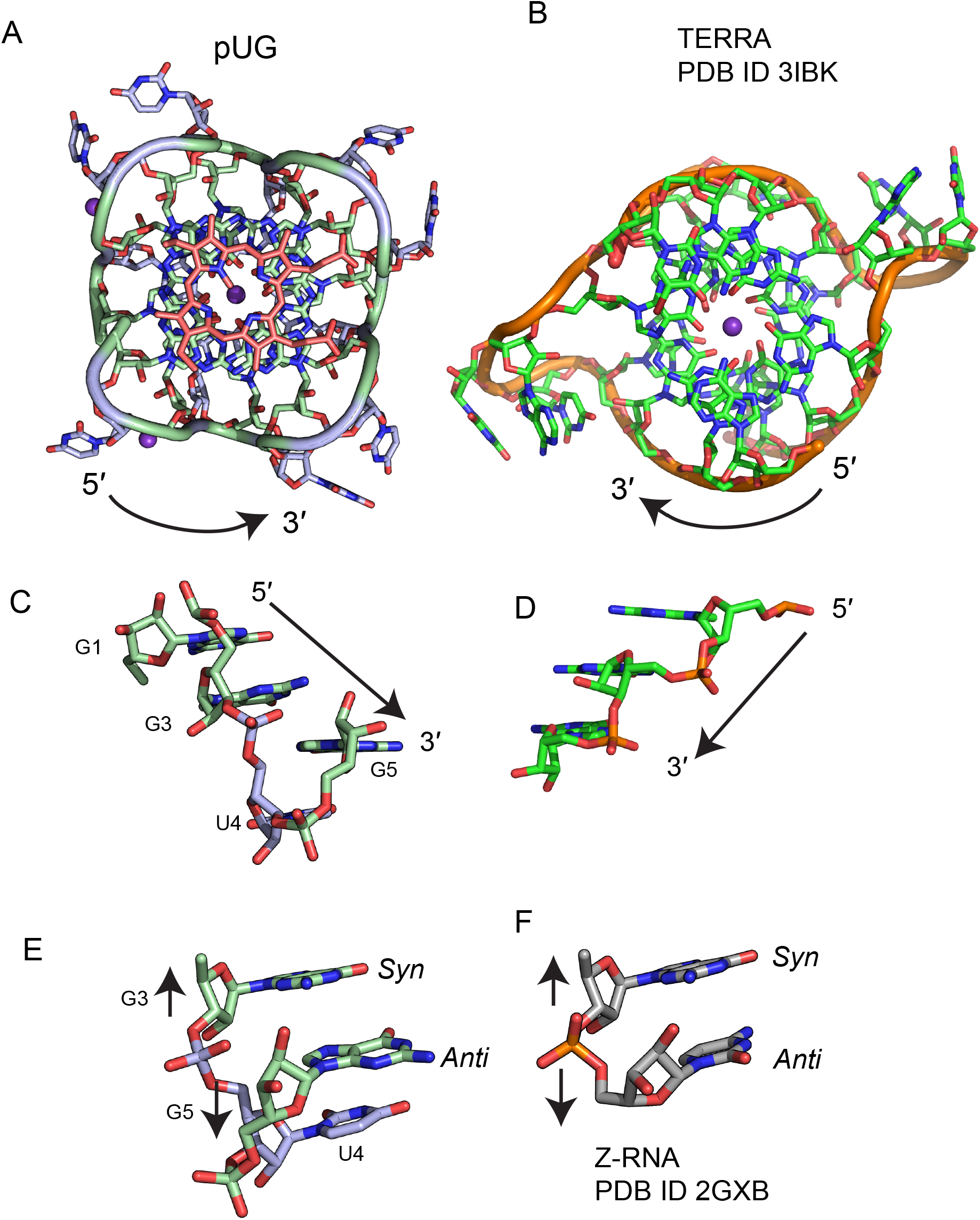
Comparison of the left-handed pUG fold (A) to the right-handed RNA G4 TERRA (PDB ID 3IBK) (B). View is looking down on the G4s with 5′ ends on top. Handedness is indicated by arrows. C. Left-handed stacking in (GU)12 vs. right-handed stacking in TERRA (D). Both views show the phosphate backbone in front of the nucleobases. E. *Syn-anti* stacking in (GU)12 with backbone inversion indicated by arrows. F. *Syn-anti* stacking in a Z-RNA duplex (PDB ID 2GXB, only one strand shown) with backbone inversion indicated by arrows.

**Supplemental Figure 8.**
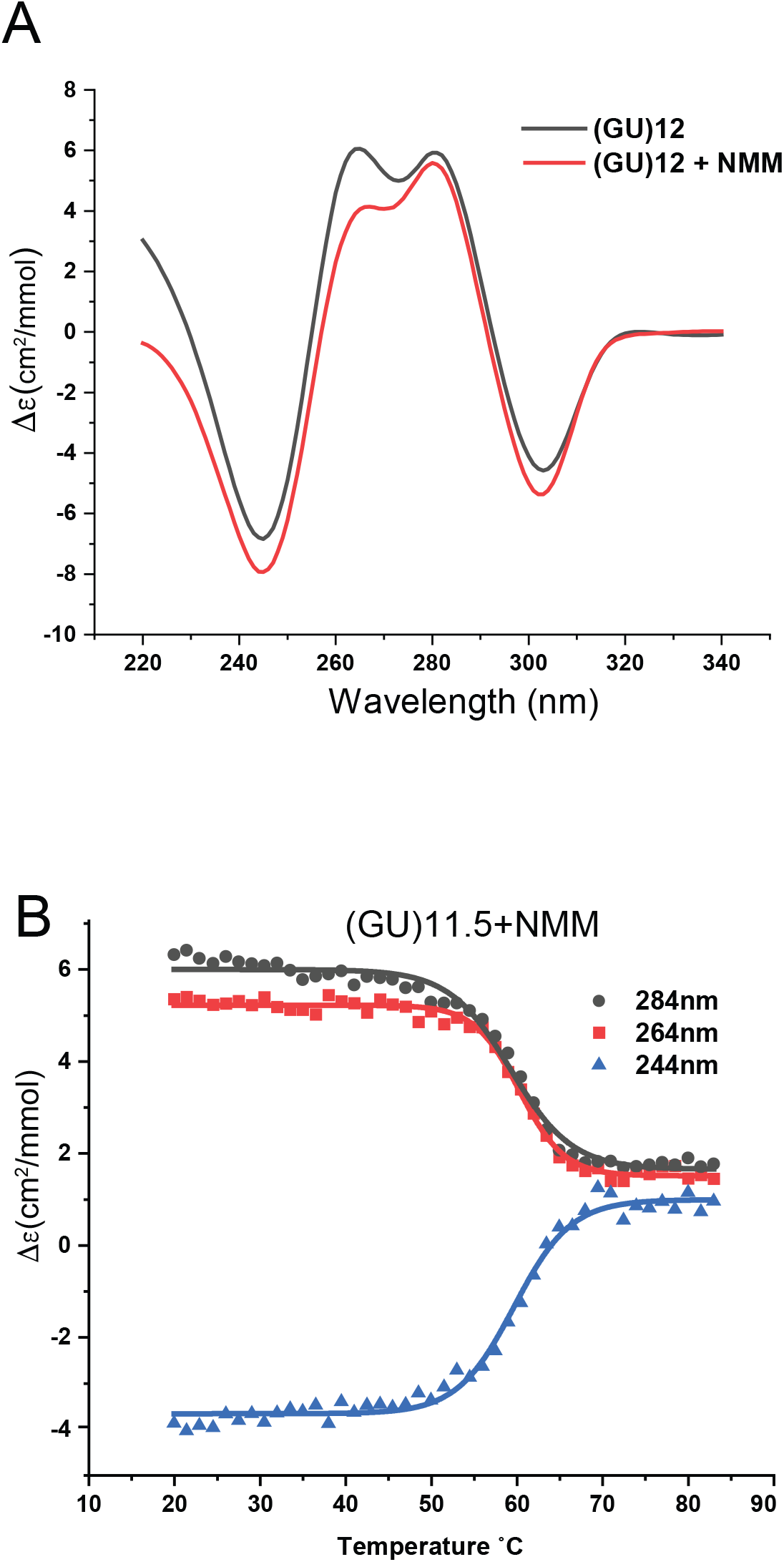
A. CD spectra of (GU)12 and the (GU)12-NMM complex. B. Thermal denaturation of (GU)12-NMM complex (1:1) monitored at three different wavelengths. The melting temperature of (GU)12-NMM is 59.7 °C.

**Supplemental Figure 9.**
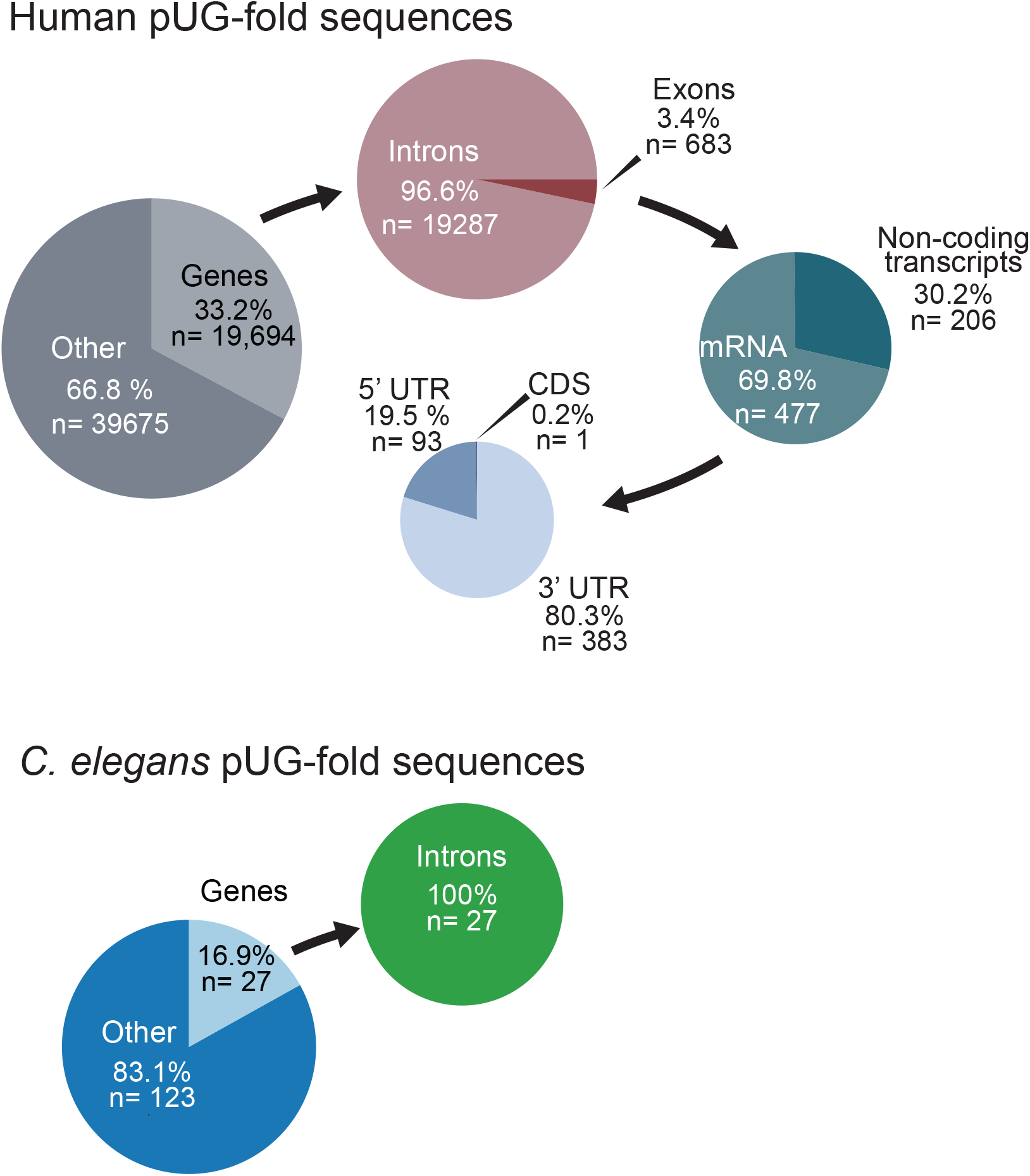
Number and distribution pUG fold sequences with 12 or more GU repeats in the human vs *C. elegans* genomes.

**Supplemental Figure 10.**
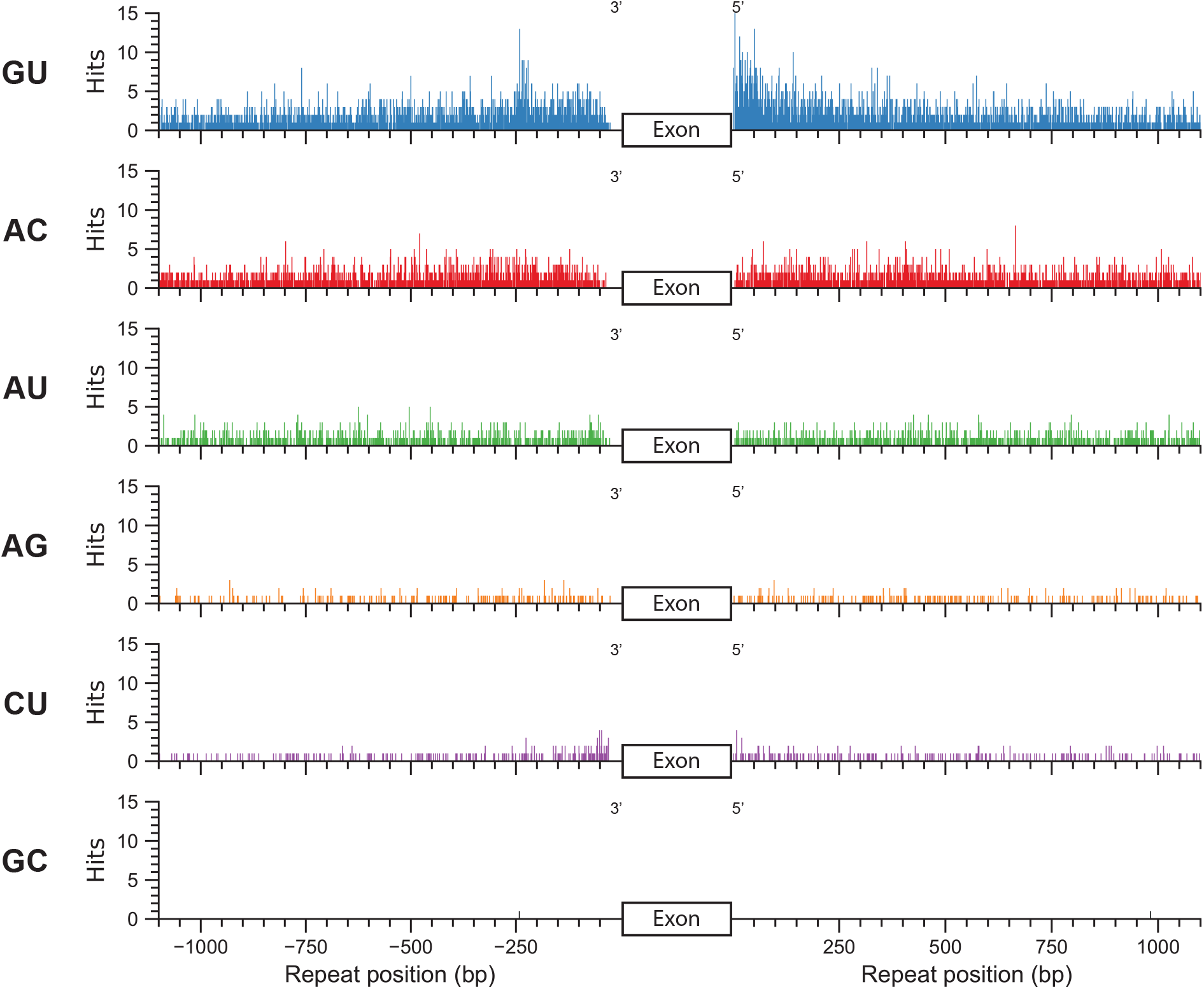
Genomic analysis of human intron sequences with dinucleotide repeat tracts of 12 or more repeats. Hits are plotted with respect to their distance from splice sites.

## Material and Methods

### RNA preparation

RNAs were either chemically synthesized from Integrated DNA Technologies or transcribed in vitro using T7 RNA polymerase. For NMR samples, ^13^C,^15^N G or U labeled RNAs were transcribed using 7.5 mM unlabeled nucleotides (Sigma) and 4 mM of ^13^C-^15^N uniformly labeled nucleotide (Silantes) in 25 mL final volume. All RNAs were purified using denaturing 15% polyacrylamide gel (8 M Urea, 89 mM Tris, 89 mM Boric acid, 1 mM EDTA) electrophoresis. The RNA was identified using UV-shadowing, cut with a razor, and removed from the gel by diffusion at room temperature overnight in 300 mM sodium acetate, 50 mM HCl, and 1 mM EDTA, pH 5.6, and then passed through a 0.2 μ filter. RNAs were further purified using a 1 or 5 ml Hi-trap Q column (GE Healthcare) that was equilibrated in (100 mM NaCl,10 mM KH_2_PO_4_, 10 mM K_2_HPO_4_, and 1 mM EDTA). RNA was washed with 20 ml of buffer and then eluted with equilibration buffer with 2 M NaCl. RNAs were concentrated in an Amicon Ultra 7 kDa filter and buffered exchanged into nuclease free water (Invitrogen), to which the indicated buffer and salt concentrations were added. For NMR samples, the 5’ triphosphate was removed with calf intestinal alkaline phosphatase (CIP) (Invitrogen) using 1U of CIP for 100 pmol of RNA in 1x of the provided CIP buffer and incubated overnight at 37 °C. CIP was removed from the samples by two rounds of phenol/chloroform extraction, followed by ethanol precipitation. RNA pellets were washed with 1 mL of cold 70% ethanol and residual ethanol was removed by speed vacuum for 10 min. NMR samples were between 0.2-0.4 mM RNA in 100 mM KCl and 20 mM potassium phosphate buffer pH 7.0 and 20 µM DSS in a volume of 300 μl. All RNA samples were pre-folded by addition of the appropriate buffer, heating the samples in 1 L of 90 °C water and slowly cooling to room temperature for 5-6 hours.

### Circular dichroism

CD RNA samples were 20 μM RNA in 20 mM Tris buffer pH 7.0 and either 150 mM KCl or 150 mM LiCl. For temperature melting experiments, RNA samples were 20 μM RNA in 20 mM potassium phosphate buffer pH 7.0 and 130 mM KCl. CD spectra were recorded in an AVIV model 420 Circular Dichroism Spectrometer using a quartz cell of 1 mm optical path length. The scans were carried at 1nm step size and 5s averaging times, measurements were taken from 210-340 nm. Spectra were measured at 25 °C with buffer subtraction and data were converted to molecular circular dichroic absorption (Δε equation 1), where θ is the raw CD signal in millidegrees, C is the RNA concentration in M, L is the cuvette pathlength in cm, and N is number of nucleotides. Thermal denaturation studies were also carried out by heating each sample between 20-85 °C with 1.5 °C intervals and 5 min. equilibration time at each temperature. Potential hysteresis was measured by comparing melting temperatures derived from experiments that ramped from low to high temperature and the reverse. Hysteresis was minimal, 3° C or less. The ellipticity was measured at 4 different wavelengths (244 nm, 264 nm, 284 nm, 304 nm) with an averaging time of 10 seconds at each temperature. The thermal unfolding profile was characterized by the millidegree signal at 244 nm, 264 nm, 284 nm to determine T_m_ values by fitting the data to the Boltzmann sigmoidal equation using Origin (Origin 2020 OriginLab corporation).

*Equation 1-molecular dichroic absorption conversion*

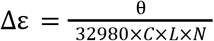

### NMM binding measurements

UV-vis absorbance spectra were collected on a Thermo Scientific NanoDrop 2000c spectrophotometer using a 1-cm polystyrene cuvette at room temperature. Titrations of NMM with (GU)_11_G were completed by stepwise additions of a 20-35 μM (GU)_11_G into a solution of 1.5-2.4 μM NMM. The pUG titrant was pre-annealed in Tris-KCl buffer (50 mM Tris, 150 mM KCl, pH 7.0) with an equivalent amount of NMM to keep the concentration of the NMM constant upon addition of (GU)_11_G throughout the titration. The titrations were terminated when consecutive additions of pUG resulted in the same spectra or when the [pUG]/[NMM] was greater than five. The spectra were measured from 220-750 nm and were deconvoluted by fitting the sum of two Gaussians centered at 378 and 397 nm corresponding to the unbound and bound NMM respectively. Complex stoichiometry was determined using a Job plot method of continuous variation following previously published protocol ^44^. Binding constants, reported as K_d_, were calculated by the direct fitting of the titration data for the bound peak according to a simple two-state, 1:1 binding model following previously published procedures ^44,45^. All data analysis was completed in GraphPad Prism software.

### NMR

All experiments were recorded on a Varian VNMRS spectrometer operating at 600 MHz (^1^H) and equipped an H/C/N cryogenically cooled probe. The temperature of the samples was regulated at 293 K throughout the experiments. The ^13^C,^15^N G-labeled sample was dissolved in H_2_O/D_2_O 90/10. A 1D ^1^H spectrum was recorded with excitation sculpting for water suppression as well as a 2D ^1^H,^15^N-HSQC optimized for G H1-N1 imino groups. A 2D ^1^H,^1^H spectrum from a NOESY ^15^N-HSQC experiment with a 200ms mixing time was used to measure NOE cross-peaks from all ^1^H signals in the molecule to the H1 imino protons of G residues. To identify hydrogen bonds, we recorded a long-range 2D ^1^H,^15^N HNN-COSY^46^ and a 2D ^1^H,^15^N HNN-TOCSY^47^ spectra. Both experiments used selective pulses on ^1^H to excite only the imino protons (IBURP2 and PC9 for excitation and REBURP for inversion). The long-range HNN-COSY was optimized to observe connectivities between H8-N7-N2 nuclei by using selective REBURP pulses on N7 and N2. The HNN-TOCSY was recorded using a 725ms long DISPI3 mixing period on nitrogen with a spinlock power of ∼300Hz to transfer magnetization between N1 imino nitrogens across hydrogen bonds. Finally, to measure RDCs for H1-N1 imino groups, 2D ARTSY^48^ spectra were recorded in isotropic conditions and after adding 20 mg/ml of PHAGE to the sample. The split of the deuterium signal for the aligned sample was stable at 26Hz throughout the experiments.

### Crystallization and structure determination

(GU)11G and (GU)12 were screened for crystallization using 200 nL RNA solution and 200 nL reservoir solution using a Mosquito (TTP Labtech, Cambridge, MA.) RNA solution consisting of 0.72 mM RNA, 0.72 mM NMM, 5 mM Tris HCl, 100 mM KCl, annealed. Optimized crystals were produced by hanging drop vapor diffusion of 1 microliter of RNA solution against a reservoir of 0.5 M Na/K tartrate, 0.1 M Tris HCl, pH 8.5, 5 mM NaOH as a pH adjuster. Crystals for both (GU)11G-NMM and (GU)12-NMM grew at 277K and were isomorphous. They were cryoprotected with reservoir solution supplemented with 30% ethylene glycol and flash-cooled by direct immersion into liquid nitrogen. Rubidium derivatized crystals were produced by soaking crystals in a solution containing 0.5 mM NMM, 0.6 m Na/Rb tartrate, 50 mM tris, pH 8.5. Na/Rb tartrate was produced by slowly adding an equimolar solution of NaOH and RbOH to solid tartaric acid, giving a clear solution. The solution was cooled overnight at 297K, yielding colorless crystals of Na/Rb tartrate. A small amount of tartaric acid was used to titrate solutions to pH 7.

Crystals were screened for diffraction at Life Sciences Collaborative Access Team (LS-CAT) and GM/CA@APS beamlines at the Advanced Photon Source (APS). Refinement and phasing data sets data were collected at (LS-CAT) beamline 21ID-D on an Eiger 9M detector. Full 360 degree sweeps of data were collected in 0.2 degree frames and reduced using XDS and autoPROC. The refinement data was collected at 1.127Å (11 keV) 170 mM sample to detector distance, 50% transmission, 50 µm beam. Phasing data was collected at 0.81413Å (15230 eV) 100% transmission, 50 µm spot, 300 mm sample to detector distance. A fluorescence scan of a similarly treated sample showed that this energy should have nearly as much anomalous diffraction as the peak at 15206 eV and be well clear of any small chemical shifts between bound and free Rb ions. Phasing data was reduced in space group P42_1_2, where the asymmetric unit could hold one quarter of the molecule. Up to four anomalous scatterers were detected with phenix.hyss in this space group, up to three on the 4-fold axis and a consistent site at a general position. The volume of the asymmetric unit in P42_1_2 can accommodate only one quarter of the RNA with reasonable packing density. It was realized that there are a series of isomorphic subgroups of P42_1_2 with progressively more voluminous asymmetric units. Reduction of symmetry to P2_1_2_1_2 with the same origin could accommodate half of the RNA and could be related to the observed diffraction pattern with two-way twinning. Further reduction of symmetry to P2_1_, with an origin shift of +/- 0.25a gives an asymmetric unit capable of holding the entire RNA and could produce the observed diffraction pattern with (nearly) perfect four-way twinning. Because the twinning was so nearly perfect, and because the twinned positions of the sites so nearly overlapped, the anomalous signal from Rb was strongest in P42_1_2, an experimental solution to the phase problem was obtained in this space group using phenix.autosolve. Using Coot were able to trace one intact RNA molecule and place one porphyrin in maps calculated in this space group. The thin-shell cross-validation set created in P42_1_2 was expanded to P2_1_. An origin shift of -0.25a placed the molecule in position for twin refinement using REFMAC5 in P2_1_ against data reduced in P2_1_.

### pUG RNA synthesis and injection

*In vitro* synthesis of pUG RNAs, preparation of injection mix, injection conditions, and data collection were performed as described previously.1 For NMM pretreatment of injected pUG RNAs, RNAs were incubated in annealing buffer (20mM Tris pH7, 100mM KCl) and 250uM NMM. RNAs were heated to 80°C, slowly cooled to 25°C (decreasing by 1°C per minute), incubated at 25°C for 12 hrs, and slowly cooled to 4°C. Mixtures were kept on ice until injection. To incorporate 7-deaza-G or 7-deaza-A into pUG RNAs, GTP or ATP was substituted with equal molar concentrations of 7-deaza-GTP (TriLink, N-1044-1) or 7-deaza-ATP (TriLink, N-1061-1) during *in vitro* transcription using MEGAscript T7 Transcription Kit (Invitrogen, AM1334). RNA was run on 6% polyacrylamide gel and stained with 0.1% toluidine blue to assess RNA integrity.

### pUG RNA chromatography

Biotinylated RNAs were synthesized by Integrated DNA Technologies (IDT). For NMM pretreatment prior to pUG RNA chromatography, biotinylated RNAs were incubated in annealing buffer (20mM Tris pH7, 100mM KCl) and 250uM NMM. RNAs were heated to 80°C, slowly cooled to 25°C (decreasing by about 1°C per minute), incubated at 25°C for 12 hrs, and slowly cooled to 4°C. Preparation of whole animal lysate, pUG RNA pull-down, gel electrophoresis, and Western blot was performed as described previously.1 To incorporate 7-deaza-G into RNAs, GTP was substituted with equal molar concentrations of 7-deaza-GTP (TriLink, N-1044-1) and synthesized *in vitro* as described in RNA preparation. 7-deaza-G RNAs and controls were preceded by a 39nt adaptor sequence (5’-GGGAGACCACGCTAGACAGCTGTTGATAATCATGTCCCC-3’). Complementary 3’ biotinylated DNA adapters (5’-ACATGATTATCAACAGCTGTCTAGCGTGGTCTCCC/3Bio/-3’) were purchased from IDT. For each experiment, 160 pmol biotinylated DNA adaptors were conjugated to 400 μg Dynabeads MyOne Streptavidin beads (Invitrogen, 65001) according to the manufacturer’s instructions. Beads were then incubated with 300 pmol *in vitro* synthesized RNAs in annealing buffer (20mM Tris pH7, 100mM KCl), heated to 80°C, and slowly cooled to 25°C (decreasing by about 1°C per minute). Beads were washed three times with annealing buffer and pUG RNA chromatography was performed as described previously.^1^

### Genome wide poly dinucleotide sequence search

A custom Python script was used to identify dinucleotide repeats in genome sequences. Linear searches of the specified repeat and its reverse complementary sequence were used to identify possible repeats in the sense and antisense DNA strands, respectively. Each poly-dinucleotide sequence was identified by its position in the genome sequence and the number of repeats. Sequence analysis was completed on the human genome (assembly GRCh38.p13) and *C. elegans* (assembly WBcel235) for 12 repeats and larger. Analysis of poly-dinucleotide sequence position in gene exons and introns was performed using human gene annotation release 109.20210226, while for *C. elegans* annotation version WS282 was used. Position analysis was performed by calculating the distance from the 5’ end of a repeat sequence to either 5’ or 3’ end of an intron or exon, whichever is closest. A positive distance was defined for a repeat closer to the 5’ end of a sequence and a negative distance defined as closer to the 3’ end of the sequence.

## Acknowledgements

Use of the Advanced Photon Source, an Office of Science User Facility operated for the U.S. Department of Energy (DOE) Office of Science by Argonne National Laboratory, was supported by the U.S. DOE under Contract No. DE-AC02-06CH11357. Use of LS-CAT was supported by NIH grant 085P1000817. Circular Dichroism data were obtained at the University of Wisconsin-Madison Biophysics Instrumentation Facility, which was established with support from the University of Wisconsin-Madison and grants BIR-9512577 (NSF) and S10RR13790 (NIH). This study made use of the National Magnetic Resonance Facility at Madison, which is supported by NIH grant P41GM136463. This study was supported by NIH grant R35 GM118131 to S.E.B.

## Author Contributions

S.R. performed CD experiments and analyzed data. J.Y. and S.G. K. performed RNA silencing experiments. E.J.M. and S.R. crystallized (GU)11.5-NMM and (GU)12-NMM complexes. C.A.B. collected diffraction data and refined the structure of the (GU)11.5-NMM and (GU)12-NMM complexes. Y.N. created the initial models for (GU)11.5-NMM and (GU)12-NMM complexes. C.E. and R.P made NMR samples and analyzed NMR data. C.E. Analyzed genomic data. R.P. Measured NMM and hemin binding to (GU)11.5. M.T. collected NMR data. E.J.M. and R.V. contributed to interpretation of structural data. M.W., S.G. K. and S.E.B. wrote the manuscript with input from all authors.

